# Statistical analyses of motion-corrupted MRI relaxometry data

**DOI:** 10.1101/2023.03.16.532911

**Authors:** Nadège Corbin, Rita Oliveira, Quentin Raynaud, Giulia Di Domenicantonio, Bogdan Draganski, Ferath Kherif, Martina F. Callaghan, Antoine Lutti

## Abstract

Consistent noise variance across data points (i.e. homoscedasticity) is required to ensure the validity of statistical analyses of MRI data conducted using linear regression methods. However, head motion leads to degradation of image quality, introducing noise heteroscedasticity into ordinary-least square analyses. The recently introduced QUIQI method restores noise homoscedasticity by means of weighted least square analyses in which the weights, specific for each dataset of an analysis, are computed from an index of motion-induced image quality degradation. QUIQI was first demonstrated in the context of brain maps of the MRI parameter R2*, which were computed from a single set of images with variable echo time. Here, we extend this framework to quantitative maps of the MRI parameters R1, R2*, and MTsat, which are computed from multiple sets of images. QUIQI allows for optimization of the noise model by using metrics quantifying heteroscedasticity and free energy. QUIQI restores homoscedasticity more effectively than insertion of an image quality index in the analysis design and yields higher sensitivity than simply removing the datasets most corrupted by head motion from the analysis. In sum, QUIQI provides an optimal approach to group-wise analyses of a range of quantitative MRI parameter maps that is robust to inherent homoscedasticity.

## 1. Introduction

The use of linear regression methods for the analysis of brain MRI data is widespread in neuroscience and neurological studies. Linear regression methods model MRI data as a linear combination of explanatory variables that may pertain to disease evolution [Beveridge et al., 2018; Ong et al., 2021; Panda et al., 2019; van der Plas et al., 2021; Scott et al., 2003], treatment [Plaikner et al., 2018], environmental factors [Hu et al., 2022; Ong et al., 2022] or phenotypes [Boots et al., 2020; Honigberg et al., 2020; Papadaki et al., 2019]. To estimate the coefficients of the combination, linear regression methods assume uncorrelated noise across measurements, sampled from normal distributions with equal variances (i.e. *homoscedasticity*). The assumption of homoscedasticity must be satisfied to ensure the validity of statistical inferences arising from the analyses of the relationship between explanatory variables and MRI data, e.g. using Student T-tests or F-tests [Hayes and Cai, 2007].

Head motion during data acquisition degrades the quality of MR images and affects brain feature estimates computed from the data [Castella et al., 2018; Esteban et al., 2017; Mortamet et al., 2009; Reuter et al., 2015; Rosen et al., 2018; Savalia et al., 2017]. In particular, motion degradation impacts the noise level of relaxometry estimates computed from raw MRI data [Castella et al., 2018]. Heterogeneous degrees of motion degradation across cohorts leads to the homoscedasticity assumption in Ordinary Least Square (OLS) analyses being violated [Lutti et al., 2022]. One solution consists of excluding the data most affected by head motion, identified from an image-based index of data quality (‘*Motion Degradation Index*’, MDI) [Castella et al., 2018; Esteban et al., 2017; Mortamet et al., 2009; Pizarro et al., 2016; Reuter et al., 2015; Rosen et al., 2018; Savalia et al., 2017]. However, determining a suitable threshold value for the index to exclude datasets of poor quality is challenging [Gilmore et al., 2021]. As a result, the resulting decrease in cohort size may lead to a sub-optimal reduction in analysis sensitivity.

The Weighted Least Square (WLS) alternative consists of including the estimate of the variance in each dataset as weights into the linear regression model. WLS analyses constitute an optimal solution because they guarantee residual homoscedasticity and preserve sensitivity. However the variance, i.e. the noise level in each individual dataset, is generally unknown. The recently introduced QUIQI method [Lutti et al., 2022] allows the variance to be estimated from the value of the MDI for each dataset through use of Restricted Maximum Likelihood (REML, [Friston et al., 2002b]). QUIQI was shown to be more effective at ensuring residual homoscedasticity and at optimizing analysis sensitivity than simple exclusion of the datasets with high MDI values.

The QUIQI method was first demonstrated for the analysis of brain maps of the transverse relaxation rate R2*, computed from a single set of raw image volumes acquired on the MRI scanner with variable echo time (TE) [Lutti et al., 2022]. R2* is primarily a correlate of iron content [Fukunaga et al., 2010; Stüber et al., 2014; Yao et al., 2009], but with sensitivity to other tissue metrics, notably myelin and water content [Bagnato et al., 2018; Hametner et al., 2018]. To disentangle the likely drivers of parameter change or difference, neuroscience studies commonly combine the analysis of multiple, complementary MRI parameters (‘*Multi-Parameter Mapping*’, [Callaghan et al., 2014; Carey et al., 2018; Schall et al., 2020]). For example, the MRI parameters R1 and MTsat show increased sensitivity to myelin content within brain tissue [Henkelman et al., 2001; Lutti et al., 2014; Sereno et al., 2013]. However, the estimation of R1 and MTsat requires several sets of raw MR images with different contrasts, and the motion level may vary across these images. Also, each MRI parameter is computed from the raw images using specific signal models [Helms et al., 2008a; Helms et al., 2008b], and is differentially sensitive to head motion as a result [Balbastre et al., 2022; Mohammadi et al., 2022]. The applicability of the QUIQI method to analyses of different types of MRI parameter maps remains to be demonstrated.

In this work, we investigate noise heteroscedasticity induced by head motion in OLS analyses of quantitative relaxometry MRI data (qMRI) computed from several sets of images. Using the global and local metrics of homoscedasticity introduced in [Lutti et al., 2022], we assess the ability of the QUIQI method to ensure the validity of statistical analyses of data degraded by motion. Multiple models of the relationship between noise estimates and indices of motion-induced image degradation are considered. We identify the optimal noise model from the global and local metrics of homoscedasticity, combined with the free energy estimates provided by REML. We compare the sensitivity of the WLS statistical analyses conducted with QUIQI with that of standard OLS analyses after exclusion of the datasets most affected by motion degradation. Finally, we compare WLS analyses conducted with QUIQI with an alternative approach based on inserting the MDI into the design matrix of OLS analyses.

## 2. Methods

### 2.1 MRI acquisition

MRI data was acquired in a large cohort of 1432 healthy research participants as part of the ‘BrainLaus’ study (https://www.colaus-psycolaus.ch/professionals/brainlaus/ [Loued-Khenissi et al., 2022; Trofimova et al., 2021]). The acquisition protocol included multi-echo T1-weighted (T1w), Proton Density-weighted (PDw) and Magnetization Transfer-weighted (MTw) scans, conducted with a custom-made 3D FLASH sequence. B1 mapping data were also acquired to correct for the effect of transmit field inhomogeneity on the qMRI maps [Lutti et al., 2010; Lutti et al., 2012]. Relevant acquisition parameters are available in [Lutti et al., 2022]. The total acquisition time was 27 minutes.

### 2.2 Map computation and processing

Computation and spatial processing of the qMRI maps were performed offline with the hMRI toolbox [Tabelow et al., 2019], as described in detail below. Image analysis was conducted using the SPM software (www.fil.ion.ucl.ac.uk/spm, Wellcome Centre for Human Neuroimaging) and customized scripts written in Matlab (R2021a, The MathWorks Inc., Natick, MA, USA).

#### 2.2.1. Computation of the qMRI maps

Maps of the MRI parameters R1, MTsat and R2* were computed from the raw T1w, PDw and MTw images of each participant. The R2* maps were computed using the ESTATICS approach from the regression of the log signal of the raw images with respect to their echo times [Weiskopf et al., 2014]. To investigate the effect of the number of raw images involved in the map calculation on homoscedasticity, maps of R2* were computed from the T1w and PDw images only (‘R2*_(2)_ maps’) and from the T1w, PDw and MTw images (‘R2*_(3)_’) (results from R2* maps computed from one set of images only (‘R2*_(1)_’) are described in [Lutti et al., 2022]).

Maps of R1 were computed from the T1w and PDw acquisitions using the rational approximation of the Ernst equation [Helms et al., 2008a]. MTsat maps were estimated using the T1w, PDw, and MTw acquisitions using a closed form expression of the MT weighted signal [Helms et al., 2008b]. In total, the performance of QUIQI was assessed on four types of quantitative maps: R2*_(2)_, R2*_(3)_, R1 and MTsat maps.

#### 2.2.2. Normalization and segmentation

Complete details of the image processing steps are available in [Lutti et al., 2022]. The MTsat maps were used for spatial normalisation of the data into the Montreal Neurological Institute (MNI) template space. The MTsat maps were segmented into maps of grey and white matter probabilities using Unified Segmentation [Ashburner and Friston, 2005]. The nonlinear diffeomorphic algorithm Dartel [Ashburner, 2007] was used for inter-subject registration of the tissue classes. To preserve the quantitative estimates, the qMRI maps were normalized into the MNI space following the voxel-based quantification procedure proposed in [Draganski et al., 2011].

#### 2.2.3. Motion Degradation Index

The MDI used for QUIQI analysis is computed by the hMRI toolbox [Tabelow et al., 2019]. This MDI was introduced in a validation study against the history of head motion that occurred during data acquisition [Castella et al., 2018]. It is calculated as the standard deviation across white matter of R2* maps computed separately from each set of multi-echo T1w, PDw and MTw images: each set of raw images was assigned a specific value of the MDI.

### 2.3 Inserting a motion degradation index into image analysis

QUIQI ensures noise homoscedasticity in analyses of MRI data affected by motion degradation by means of weighted least square analyses (WLS). The weights, specific to each dataset of an analysis, are computed from a noise covariance matrix (*V*) modelled as a linear combination of matrices built from the MDI values of the analysis data (‘basis functions’).

From the heuristic analysis of the empirical relationship between analysis residuals and the MDI [Lutti et al., 2022], these basis functions were set to contain powers of the MDI values:

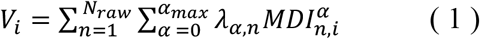

where *V*_*i*_ is the i^th^ diagonal element of the covariance matrix *V*, i.e. the noise estimate of the i^th^ dataset. 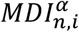 is the *α*^*th*^ power of the MDI of the i^th^ dataset and the n^th^ contrast weighting. The basis functions and the resulting noise covariance matrices *V* were diagonal because noise is uncorrelated between datasets (i.e. participants).Estimation of *V* involved the MDI of each raw image involved in the computation of the qMRI data to be analysed (index *n* in eq.1), and separate basis functions were computed from each MDI. *Nraw* is the number of raw images involved in this computation: *Nraw* =2 for R2*_(2)_ and R1 maps as these data were estimated from the PDw and T1w raw images. *Nraw*=3 for MTsat and R2*_(3)_ maps as these data were estimated from the PDw, T1w and MTw raw images (see section 2.2.1).

The optimal value of *α*_*max*_, the maximum power of the MDI used to compute the noise covariance matrix, was determined in a model comparison (see section 2.5.1).

*V* and *λ*_*α,n*_ were estimated using the Restricted Maximum Likelihood (REML) algorithm as implemented in SPM12 [Friston et al., 2002a]. The subsequent estimation of the weights 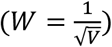 for WLS analyses was conducted with the standard analysis tools of SPM12.

### 2.4 Measures of analysis validity

The results of OLS and WLS analyses were compared using local and global metrics of heteroscedasticity, and a measure of free energy. These metrics were also used to identify the optimal model of the dependence of the noise in qMRI maps on image quality.

#### 2.4.1 Residual heteroscedasticity

We characterized noise heteroscedasticity from the maps of the residuals *ϵ*, computed for each participant and qMRI maps after estimation of the coefficients of the linear model.

##### Global heteroscedasticity

We computed a global noise index *var*(*ϵ*), as the spatial variance of the residual maps across a whole tissue type (i.e. grey or white matter) [Lutti et al., 2022]. Consistent with Eq 1., the effect of motion degradation on the global noise index was modelled as a cubic polynomial combination of the MDI of all the raw images involved in the computation of the analysis data. For R2*_(2)_ and R1 maps, the MDI of the T1w and PDw raw images were considered. For R2*_(3)_ and MTsat maps, the MDI of the T1w, PDw and MTw raw images were considered. Fitting the global noise index with the assumed polynomial combination of the MDI led to the estimation of *var*(*ϵ*)_*fit*_, the modelled dependence of the global noise level on motion degradation.

The global heteroscedasticity index was taken as the coefficient of determination 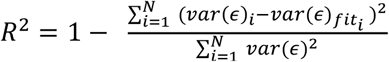. A high value of R^2^ (∼1) indicates a strong dependence of the global noise index on motion degradation and high heteroscedasticity. Conversely, a low value of R^2^ (∼0) indicates a weak dependence of the global noise index on motion degradation and low noise heteroscedasticity.

A graphical rendering of heteroscedasticity is achieved by plotting *var*(*ϵ*) against *var*(*ϵ*)_*fit*_ : a linear relationship between *var*(*ϵ*) and *var*(*ϵ*)_*fit*_ reflects high heteroscedasticity. The absence of relationship between *var*(*ϵ*) and *var*(*ϵ*)_*fit*_ reflects low heteroscedasticity.

##### Local heteroscedasticity

At the local scale, we assessed noise heteroscedasticity in each image voxel using the Engle’s Arch test [Engle, 1982], which tests for no linear relationship between consecutive samples of a series of squared residuals. At a given voxel, the residual series was computed from the residual maps at this voxel location. The series samples were organized according to the predicted impact of motion degradation by arranging them in ascending order of the polynomial combination of MDIs (*var*(*ϵ*)_*fit*_), associated to the tissue class of the given voxel.

The Engle’s Arch test was conducted at each voxel of the residual maps, with a maximum lag of 40 points. The local index of heteroscedasticity was the proportion of voxels with significant heteroscedasticity (i.e. rejecting the null hypothesis), calculated after false discovery rate correction using the Benjamini-Hochberg procedure (p<0.05; [Glickman et al., 2014]).

#### 2.4.2 Free energy

REML estimates the hyperparameters λ that are used to compute the matrix *V* (Eq.1) by maximizing the evidence lower bound objective (ELBO) function, a measure of negative variational free energy. Intuitively, the ELBO favours the accuracy and penalizes the complexity of the model. We used the ELBO estimates provided by the REML implementation in SPM12 [Friston et al., 2002a] to identify the optimal model of the relationship between noise in the qMRI maps and the degradation of the raw images induced by head motion (see section 2.5.1).

### 2.5 Image analyses

Linear regression analysis was conducted on qMRI maps of R2*_(2)_, R2*_(3)_, R1 and MTsat. This allowed the assessment of noise heteroscedasticity in WLS and OLS analyses, separately for different MRI parameters and different numbers of raw image types involved in their estimation. The quantitative MRI data were modelled as the linear combination of 5 regressors that represented age, square and cubic values of age, gender and brain volumes.

#### 2.5.1 Noise model comparison

The coefficients of the general linear model were estimated from the full dataset (N=1432). OLS analyses were conducted with *α*_*max*_ = 0 in Eq.1, i.e. assuming uniform noise level across all datasets as would be done in a standard analysis. WLS analyses were conducted with 5 models of the noise covariance matrix: *α*_*max*_ =2 to 5 with *λ* ∈ ℝ, and *α*_*max*_ =4 with *λ* ∈ ℝ^+^. The latter model was considered to assess the effect of enforcing positivity of the λ hyperparameters on noise heteroscedasticity and free energy, and for consistency with [Lutti et al., 2022].

OLS and WLS analyses were compared by considering differences in heteroscedasticity and free energy. Across the WLS analyses, the optimal noise model was identified as that which led to the lowest values of heteroscedasticity and the largest increase in ELBO.

#### 2.5.2 Age sensitivity

We compared the sensitivity of WLS analyses against that of standard OLS analyses using statistical F-tests of age-related differences in R2*_(2)_, R2*_(3)_, R_1,_ and MTsat. The analyses were performed on a subsample of the full dataset, obtained by randomly selecting up to 10 images per age bin of 5 years. The size of the resulting dataset was 123, close to the typical cohort size of similar studies [Callaghan et al., 2014].

The WLS analyses were conducted with the optimal noise model (see section 2.5.1). OLS analyses were conducted after exclusion of the datasets most degraded by head motion, i.e. with the highest MDI values averaged across the raw images involved in the calculation of the qMRI maps. Exclusion of 3, 7, 13, 20 and 30% of the most degraded datasets was considered [Castella et al., 2018; Esteban et al., 2017; Mortamet et al., 2009; Pizarro et al., 2016; Reuter et al., 2015; Rosen et al., 2018; Savalia et al., 2017]. The optimal fraction of excluded datasets was identified as the smallest value leading to local and global heteroscedasticity estimates comparable to those obtained in WLS analyses with the optimal noise model.

#### 2.5.3 Specificity

To assess the specificity of the OLS and WLS analyses, we recorded the rate of false positives in two types of analyses frequently conducted in neuroscience studies. I) In a subset of the full dataset with up to 10 images per age bin of 5 years (N = 123), the participants’ age was randomly assigned from a uniform distribution ranging from the minimum to the maximum age of the data subsets and statistical F-tests of age-related differences in R2*_(2)_, R2*_(3)_, R_1_ and MTsat maps. II) In a subset of the full dataset within a narrow age range (56–58 y.o.; N = 129), we conducted unbalanced comparisons of a group of 10 qMRI maps with a group of 119 maps, using two-sample T-tests. Given that no age-related effect would be expected in these analyses, any significant effects were deemed to be false positives.

We conducted analyses I and II 1,000 times for different subsets of data and monitored the rate of significant results, i.e. false positives, across repetitions at the cluster level (p < 0.05, FWE-corrected), with a cluster forming threshold of p < 0.001 uncorrected.

#### 2.5.4 Inserting the Motion Degradation Index in the analysis design

In a subset of data in a narrow age range (56–58 y.o.; N = 129), we considered an alternative method to QUIQI to correct motion degradation effects, which consists of inserting the MDI as a confounding factor in the analysis design. Here, powers of 1 to 4 of the MDI values were included as regressors in the design matrix. We compared noise heteroscedasticity between OLS analyses and WLS analyses with the optimal noise model (*α*_*max*_ = 4, *λ* ∈ ℝ, identified from 2.5.1).

## 3. Results

### 3.1 Noise model comparison

With standard OLS analyses, global heteroscedasticity ranges from 0.44 to 0.7 across the different types of analysed qMRI data, and the fraction of voxels with significant local heteroscedasticity ranges from 0.42 to 0.85 (Fig. 1). Global and local heteroscedasticity are largely comparable across the analyses of R2*_(2)_ and R2*_(3)_ maps, suggesting little effect of the number of raw images involved in the computation of the qMRI maps on heteroscedasticity. The generally lower level of heteroscedasticity in analyses of R1 and MTsat maps suggests a bigger effect of the type of analysed qMRI data.

**Figure 1:**
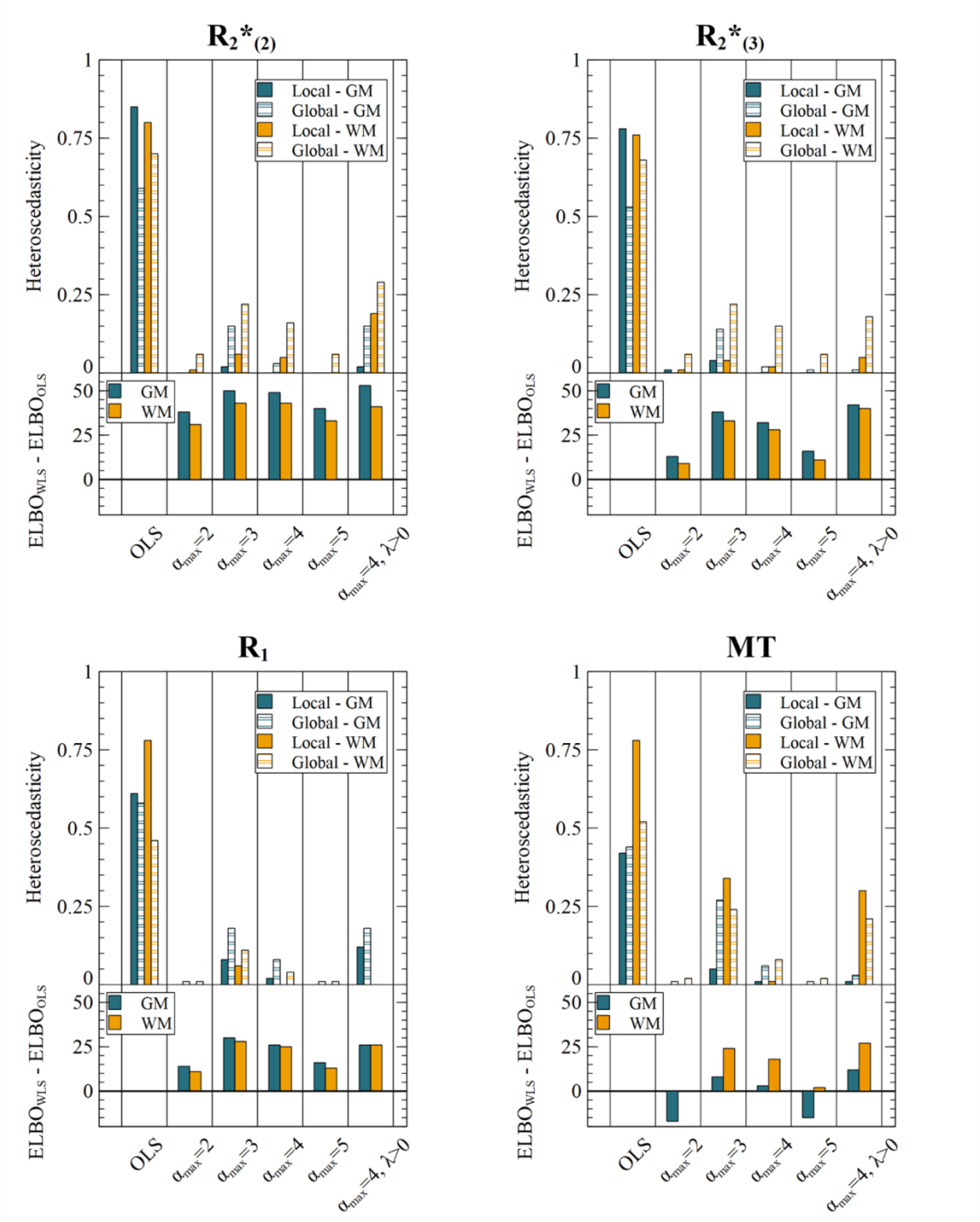
WLS analyses reduce heteroscedasticity and increase ELBO. Each graph shows heteroscedasticity and ELBO levels for different qMRI maps R2*_(2)_ (top left), R2*_(3)_ (top right), R1 (bottom left) and MTsat (bottom right)). The top part of each graph shows local and global heteroscedasticity levels for OLS analyses and WLS analyses with different noise models (i.e. *α*_*max*_), in white matter (WM) and grey matter (GM). The bottom part shows differences in ELBO between WLS and OLS analyses. WLS analyses strongly reduce noise heteroscedasticity, for all noise models and qMRI maps. WLS analyses also lead to an increase in ELBO, except for two noise models in the GM of MTsat maps.

WLS analyses reduce global and local heteroscedasticity for all types of qMRI data, models of covariance matrix and tissue types (Fig.1). Local heteroscedasticity is below 0.05 with *α*_*max*_ = 2.4 or 5, *λ* ∈ ℝ). With these models, the global measure R^2^ is below 0.16. The two alternative models (*α*_*max*_ = 3, *λ* ∈ ℝ and *α*_*max*_ = 4, *λ* ∈ ℝ^+^) lead to higher global and local heteroscedasticity, particularly for MTsat data.

Across the noise models with (*λ* ∈ ℝ), the increase in ELBO compared to OLS analyses reach a maximum for *α*_*max*_ = 3, closely followed by *α*_*max*_ = 4. Noise models with *α*_*max*_ = 2 and *α*_*max*_ = 5 lead to a decrease in ELBO in analyses of MTsat maps. The gains in ELBO compared to OLS analyses are higher in analyses of R2*_(2)_ data followed by R2*_(3)_, R1, and MTsat data, suggesting an effect of both the type of analysis data and of the number of raw image volumes involved in their computation.

Enforcing positivity for the REML hyperparameter (i.e. *λ* ∈ ℝ^+^) further increased the ELBO. However, noise heteroscedasticity is comparably high for this model of the noise covariance.

Noise homoscedasticity is essential to ensure the validity of statistical inference and was the key requirement in our selection of the optimal model of the noise covariance. Among the models that ensure noise homoscedasticity (*α*_*max*_ = 2.4 or 5, *λ* ∈ ℝ), we deemed *α*_*max*_ = 4, *λ* ∈ ℝ to be optimal as it leads to the highest increase in ELBO compared to OLS analyses, for all types of analysis data and both tissue types. With this model, analysis residuals depend only weakly on the value of the MDI of the analyses data (i.e. global heteroscedasticity R^2^ ≤ 0.16 see Fig.2A). The ELBO systematically increases compared to OLS analyses, by an amount that varies according to the type of qMRI data and number of raw image volumes involved in the map calculation (see Fig.2B).

**Figure 2:**
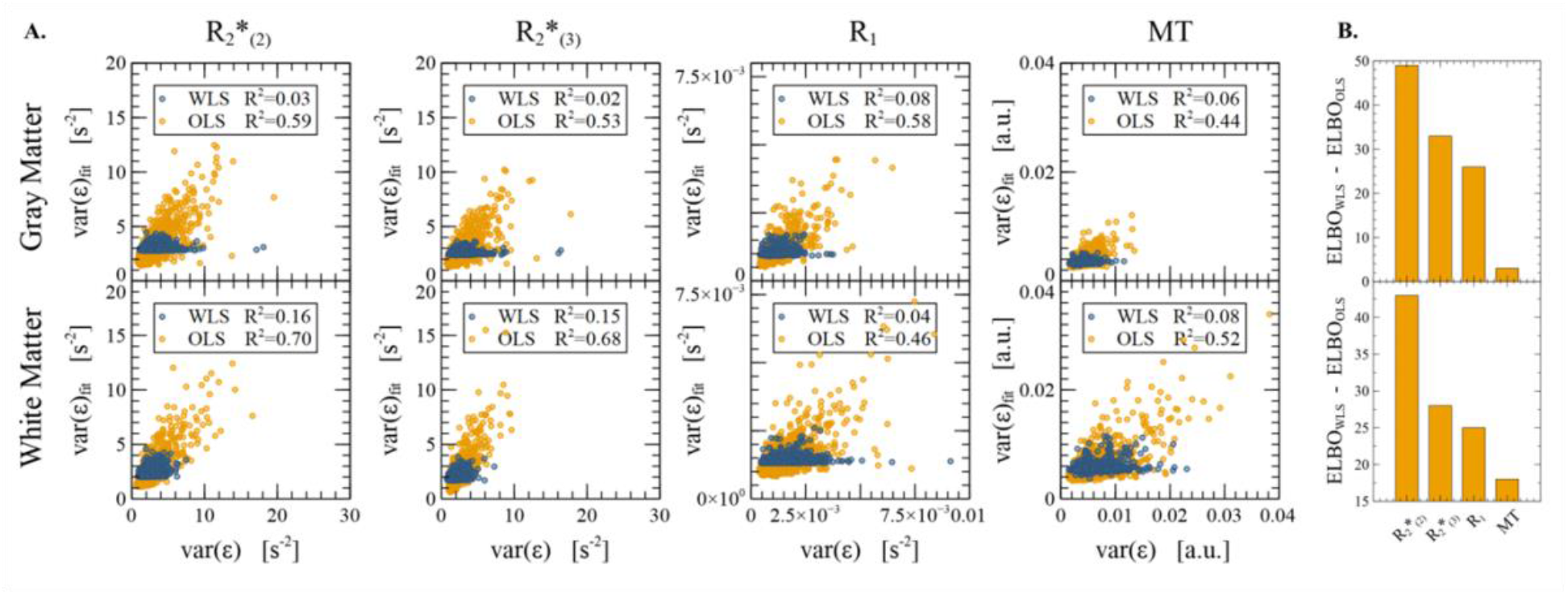
Noise homoscedasticity and ELBO increase with the optimal noise model. (a) With the optimal noise model (*α*_*max*_ = 4, *λ* ∈ ℝ), global heteroscedasticity (R^2^) does not exceed 0.16, for all types of qMRI maps and in both grey and white matter. (b) WLS analyses lead to an increase in ELBO, for all types of qMRI maps and in both grey and white matter. This increase varies according to the type of qMRI data and the number of raw image volumes involved in the map calculation.

### 3.2 Age sensitivity

Differences in age sensitivity between OLS and WLS analyses conducted on the full dataset show substantial effects, both positive and negative (Supporting Figure S1). This is consistent with the effect of noise heteroscedasticity in OLS analyses, which might lead to under- or over-estimation of the noise level - both of which invalidate associated inferences. To ensure a valid comparison of age sensitivity with WLS analyses, we investigated the exclusion from the OLS analyses of the datasets most affected by head motion, which we identified from their high MDI values. Overall, global and local noise heteroscedasticity decrease with increasing fraction of excluded datasets (Fig.3). After exclusion of the 30% of the datasets with the highest MDIs, global heteroscedasticity lies in the same range as WLS analyses overall (R^2^∼0.1-0.2), except for R2*_(2)_ and R2*_(3)_ data in white matter (R^2^=0.34). However, local heteroscedasticity remains generally higher than for WLS analyses (<0.05), and reaches 0.12 in R2* data and 0.25 in the white matter of MTsat data (solid bars in Fig.3). Higher exclusion fractions, which would have reduced local heteroscedasticity further, were deemed too prohibitive to be investigated.

**Figure 3:**
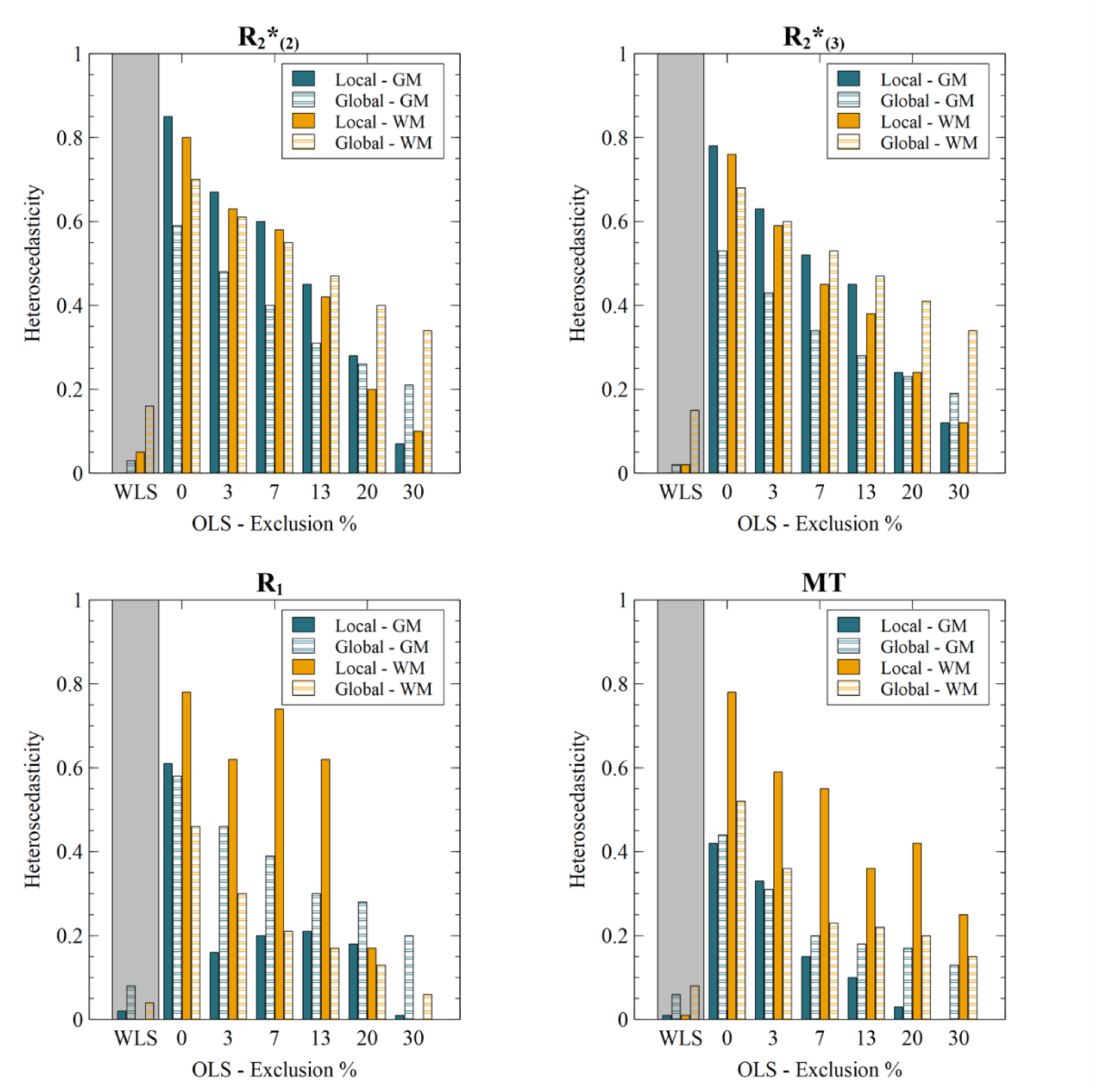
Exclusion of the most degraded datasets improves noise heteroscedasticity in OLS analyses. After exclusion of 30% of the datasets, global heteroscedasticity lies in the same range as WLS analyses overall (R^2^∼0.1-0.2), except for R2*_(2)_ and R2*_(3)_ maps in white matter (R^2^=0.34). However local heteroscedasticity remains generally higher than for WLS analyses (<0.05), and reaches 0.12 in R2* data and 0.25 in the white matter of MTsat data.

The age sensitivity of the WLS analyses was compared with that of OLS analyses after exclusion of the 30% of datasets with the highest MDIs. Statistical F-maps of age-related changes in qMRI data, obtained from WLS analyses, exhibit predominant features that are consistent with previous findings from the literature (Fig.4, [Callaghan et al., 2014]). These include an increase in R2* with age in sub-cortical areas attributed to a local increase in iron concentration, and a decrease in MTsat and R1 in frontal white matter attributed to fibre demyelination. With WLS analyses, statistical scores are larger and the significance threshold is lower due to the higher number of samples (see inset in Fig. 4). The number of voxels above significance increases by a factor 2.8 to 4.3 with WLS analyses, except for the R1 and MTsat parameters in white matter, where the increase is of 20% and 5% respectively (Fig. 5). With WLS analyses, the spatial distribution of significant voxels shows enhanced symmetry between the left and right hemispheres (Fig. 5). Regions of significant voxels also show improved spatial continuity.

**Figure 4:**
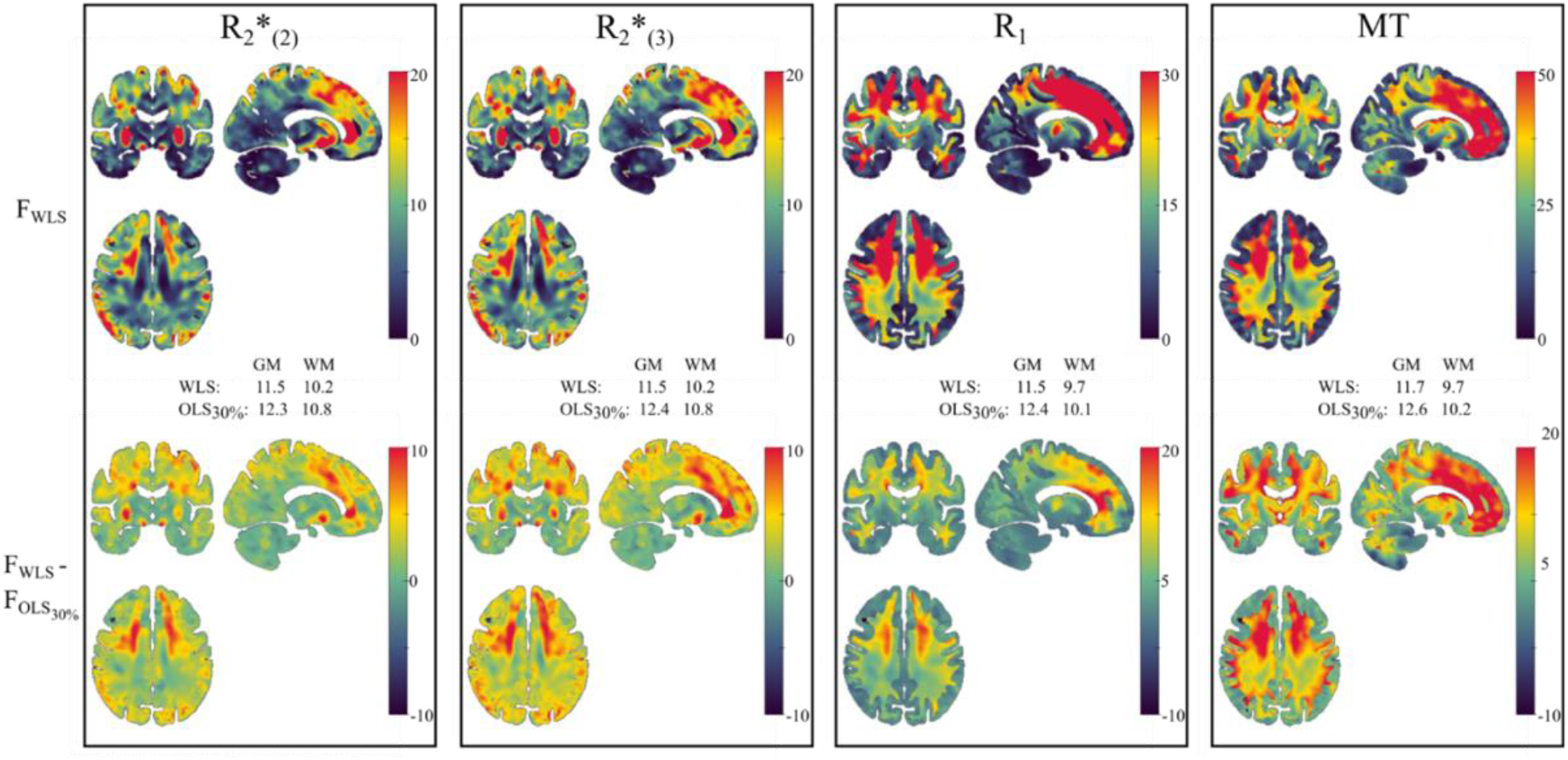
WLS analyses show increased sensitivity to brain differences. Statistical F-maps of age-related changes (F_WLS_), obtained in WLS analyses of 123 samples of the full dataset, show predominant features that include an increase in R2* with age in sub-cortical areas and a decrease in MTsat and R1 in frontal white matter (top). WLS analyses exhibit higher Fscores than OLS analyses after exclusion of 30% of the most corrupted data, in all types of qMRI data and throughout grey and white matter (bottom). For each type of qMRI maps, the inset indicates the threshold for statistical significance (p<0.05, FWE corrected).

**Figure 5:**
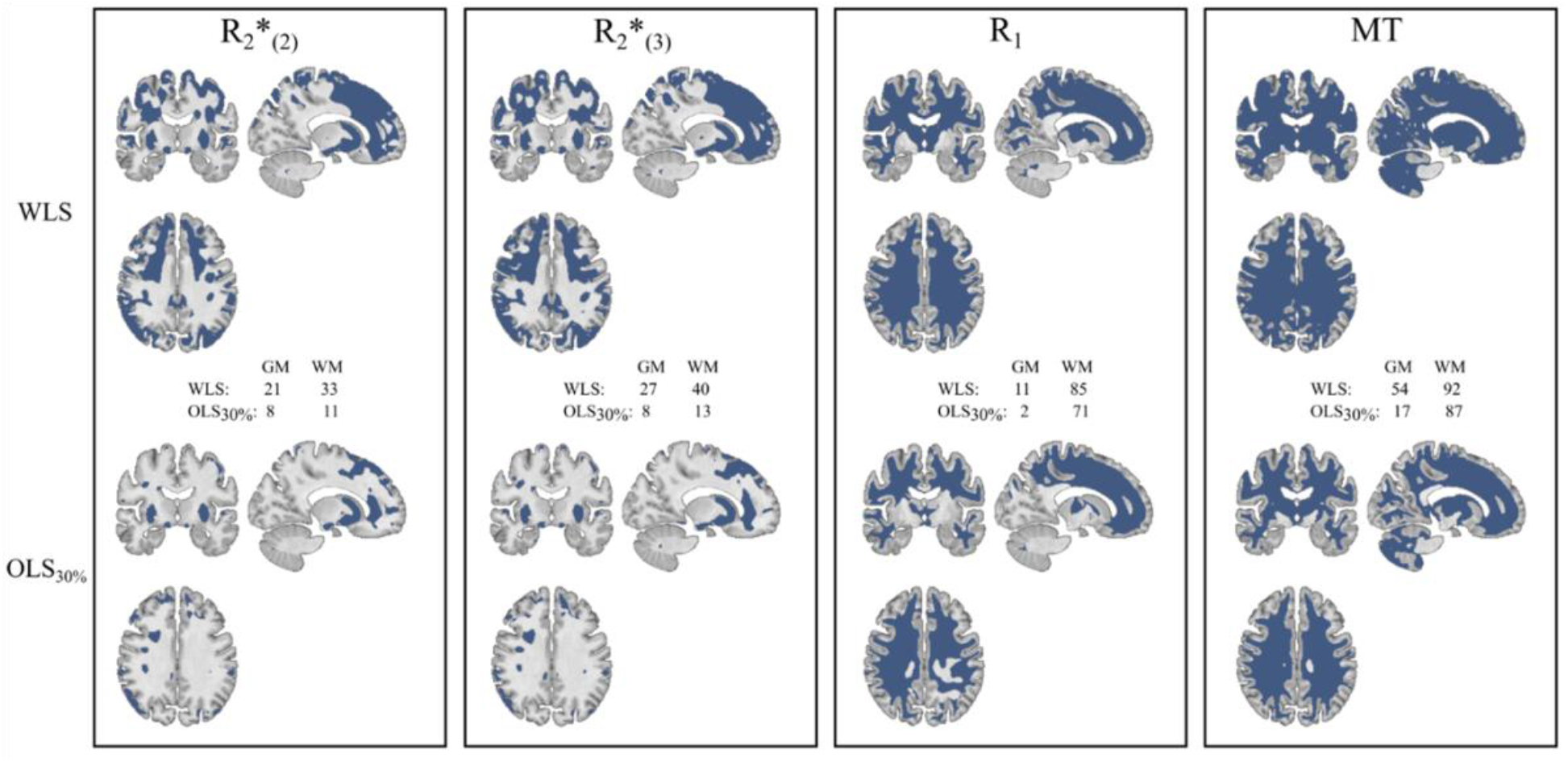
WLS analyses increase the spread of significant voxels in statistical maps. Maps of voxels with significant age-related changes (blue) (p<0.05, FWE corrected) show increased spatial extent with WLS analyses (top) than OLS analyses with exclusion of 30% of the most corrupted data (bottom), for all types of qMRI data. The insets indicate the fraction of significant voxels in grey and white matter.

### 3.3 Specificity

OLS analyses and WLS analyses with the optimal noise model (*α*_*max*_ = 4, *λ* ∈ ℝ) show equivalent rates of false positives in both F-tests and imbalanced two-sample T-tests (Table 1). On average, OLS and WLS analyses show significant results at the cluster level in 5.4% and 6.5% of the tests, in agreement with the expected rate of 5%. The rates of false positives are slightly lower for the tests with shuffled age regressor but remain comparable between OLS and WLS analyses (3.1% and 2.5% respectively).

**Table 1 :**
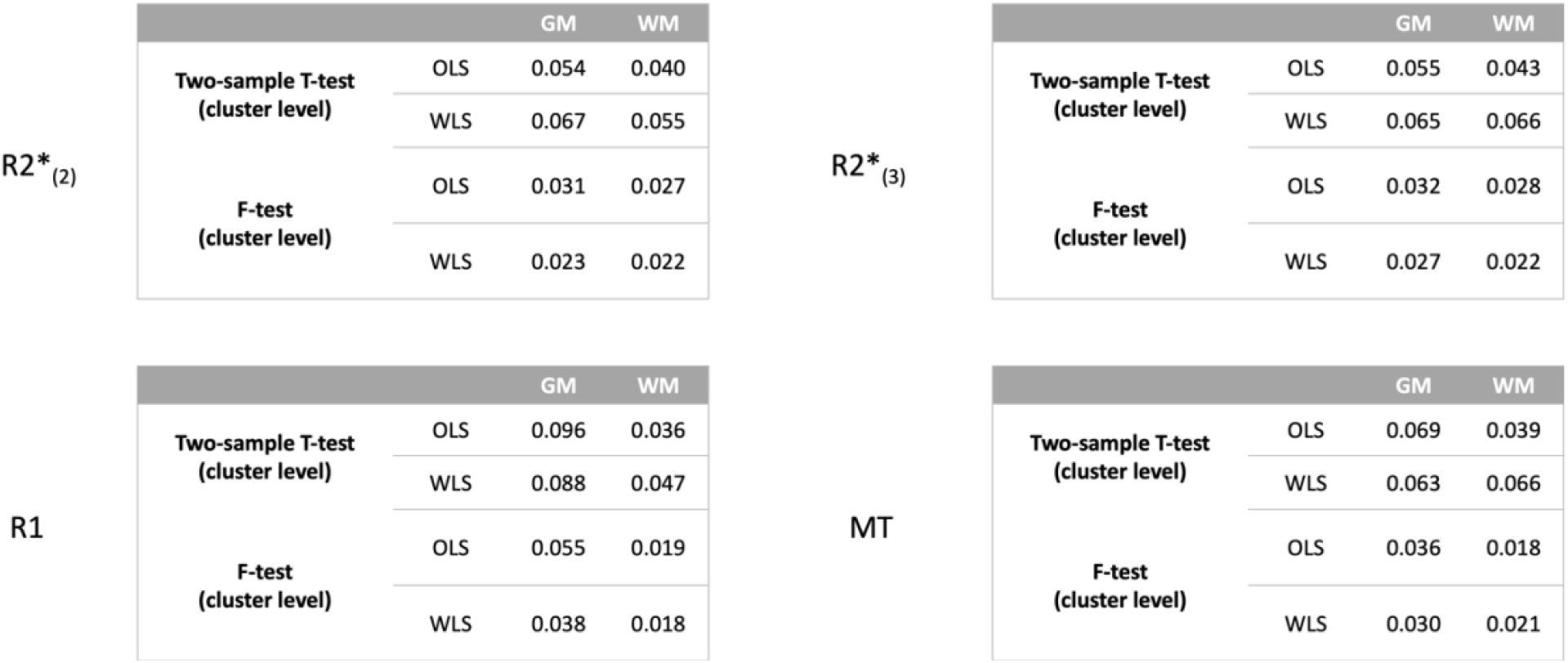
Specificity is preserved with WLS analyses. Specificity of WLS and OLS analyses of R2*(2), R2*(3), R1 and MTsat data, in grey and white matter. The rate of false positive clusters (p < 0.05) were obtained with two tests: 1/ Group comparison of two sub-groups within a narrow age range with a two-sampled T-test 2/ Age-associated differences after random assignment of the participants’ age with an F-Test. Any positive result is a false positive. The cluster forming threshold was p < 0.001 uncorrected.

### 3.4 Inserting the Motion Degradation Index in the analysis design

The level of noise heteroscedasticity in WLS analyses conducted with QUIQI was compared with that of OLS analyses after insertion of the MDI into the design matrix. Residual heteroscedasticity remains present in OLS analyses (R^2^_WLS_< R^2^_OLS_), for all data and tissue types (Fig. 6).

**Figure 6:**
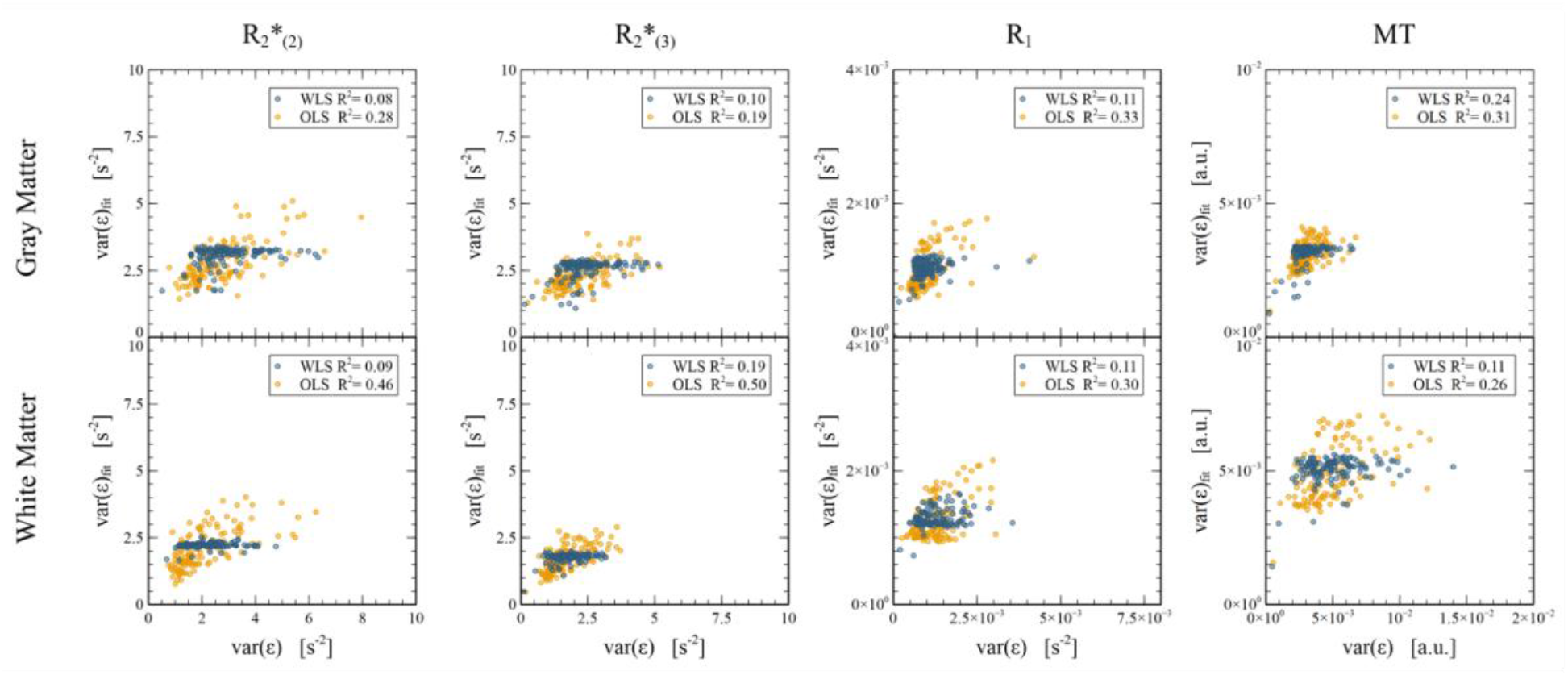
Including the motion degradation index as regressors in the design matrix does not restore homoscedasticity. Despite inserting the MDI in the design matrix, OLS analyses exhibit a high level of global noise heteroscedasticity (high R^2^), for all types of qMRI maps and in both grey and white matter. Homoscedasticity is restored with WLS analyses.

## 4. Discussion

Analyses of MRI data using linear regression rely on the assumption of noise homoscedasticity to ensure the validity of statistical inference. Degradation of MRI data quality due to head motion invalidates this assumption. The resulting mis-estimation of the noise variance leads to erroneous statistical inference and increased risks of false positives or negatives. The QUIQI method restores the validity of statistical analyses by conducting weighted least-square estimations, with weights that are computed from an index of data degradation due to head motion. The benefit of QUIQI has recently been demonstrated from a large dataset of R2* maps computed from a single set of raw images [Lutti et al., 2022]. Here, we extended this method to brain maps of the MRI parameters R2*, R1 and MTsat, computed from multiple sets of raw images. The impact of restoring statistical validity is substantial and consistent with under- or over-estimation of the noise variance in OLS analyses (Supporting Figure S1). WLS analyses show higher sensitivity to brain differences across a dataset than OLS analyses that excluded the most corrupted data (Fig. 4-5). WLS analyses are also more effective at ensuring homoscedasticity than insertion of the MDIs as regressors in the design matrix of the analysis.

QUIQI requires modelling of the noise covariance matrix from a Motion Degradation Index (MDI) of the analysis data. The relationship between analysis residuals and the MDI in an OLS analysis constitutes a good baseline to infer a suitable form for this model [Lutti et al., 2022]. For the current application, we chose to base this model on polynomial combinations of the MDI of each of the raw images involved in the computation of the R2*, R1 and MTsat maps. We compared candidate models, with different polynomial degrees and constraints on the hyperparameter *λ*, from measures of local and global noise heteroscedasticity, and from ELBO - a measure of free energy. WLS analyses strongly reduced heteroscedasticity for all the types of qMRI data analysed here. Also, the same subset of models consistently led to optimal restoration of homoscedasticity. From this subset, we identified the optimal noise model that led to the highest increase in ELBO compared to OLS analyses. The use of higher powers of the MDI increased the complexity of the noise covariance model, leading to a decrease in the ELBO.

The sensitivity of OLS analyses and WLS analyses conducted with QUIQI were compared in the context of age-related effects [MacDonald and Pike, 2021]. For our comparison, the 30% of datasets with the highest MDI were removed from the OLS analyses to achieve a similar level of homoscedasticity than the WLS analyses. The higher sensitivity of the WLS analyses conducted with QUIQI arises from its ability to exploit the full dataset to increase statistical power. With WLS analyses, the spatial distribution of significant voxels shows improved biological plausibility in the form of greater spatial continuity and enhanced symmetry between the left and right hemispheres. This increase in sensitivity preserves specificity, i.e. the rate of false positives in the statistical analyses remained under control.

QUIQI is available for use within the hMRI toolbox (https://hmri-group.github.io/hMRI-toolbox/). This implementation relies on the REML algorithm of SPM12 that is commonly used to correct for temporal correlations and group-level analyses of fMRI data [Friston et al., 2002a]. Following specification of the design matrix (‘*factorial design*’), the *QUIQI_Build* module builds a dictionary of basis functions from MDI values provided by the user. By default, *QUIQI_Build* computes the basis functions as powers of the MDI that are specified by the user. Alternatively, the noise model can be readily adapted to different analysis datasets and MDIs (see hmri_quiqi_build.m). By default, no constraint is imposed on the hyperparameters (see [Lutti et al., 2022] for details on adding a positivity constraint). The *QUIQI_Check* module can be used after image analysis to estimate global heteroscedasticity in the data. Here, the degree of the polynomial used to estimate var(ε)_fit_ is independent of the powers of the MDI used to model the noise covariance matrix in *QUIQI Build*.

## 5. Conclusion

Degradation of image quality due to head motion invalidates the homoscedasticity assumption in the statistical analysis of MRI data. Here, we extended the QUIQI method to the analysis of brain maps of the MRI parameters R2*, R1 and MTsat, computed from multiple sets of raw images. QUIQI restores homoscedasticity and the validity of statistical inference, and allows for optimization of the noise model using specially-dedicated metrics of heteroscedasticity and free energy. QUIQI is more effective at ensuring homoscedasticity than regressing out the image quality indices, and yields higher sensitivity than removal of the datasets most corrupted by head motion from the analysis. QUIQI is available in the hMRI toolbox for the study of brain differences from MRI data.

---

**Figure S1:**
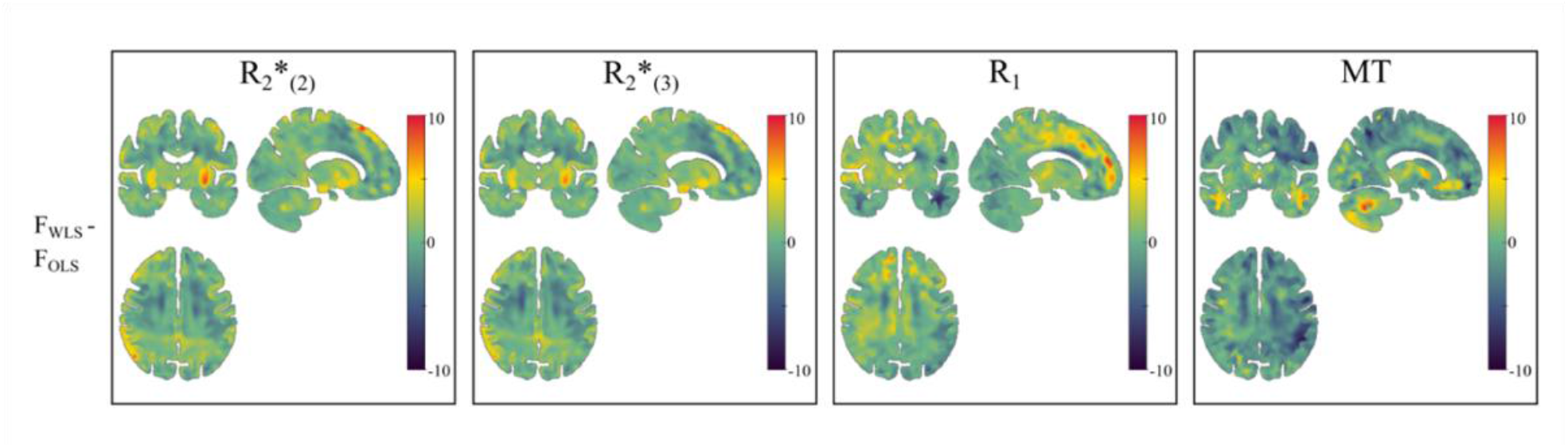
Noise heteroscedasticity leads to under- or over-estimated statistical results. Differences in statistical F-maps between OLS and WLS analyses conducted on the full dataset are substantial and may be positive or negative, consistent with the effect of noise heteroscedasticity in OLS analyses.

## Notes

**Funding statement** This work was supported by the Swiss National Science Foundation (grant no 320030_184784 (AL)), the Partridge foundation and the ROGER DE SPOELBERCH Foundation. The Wellcome Centre for Human Neuroimaging is supported by core funding from the Wellcome [203147/Z/16/Z]. BD is supported by the Swiss National Science Foundation (NCCR Synapsy, project grant numbers 32003B_135679, 32003B_159780, 324730_192755 and CRSK-3_190185) and the Fondation Leenaards. The MRI data were acquired on the MRI platform of the Clinical Neuroscience Department, Lausanne University Hospital.

**Conflict of interests disclosure** The authors declare no potential conflict of interest

https://zenodo.org/record/7612032#.Y-OwlXbMKUl

https://hmri-group.github.io/hMRI-toolbox/

https://zenodo.org/record/7692074#.ZACk9HaZOUk

### Competing Interest Statement

The authors have declared no competing interest.

### Summary of Updates

Details on journal submisssion removed

