## Supplementary material for "Statistical analyses of motion-corrupted MRI relaxometry data": Fig S1

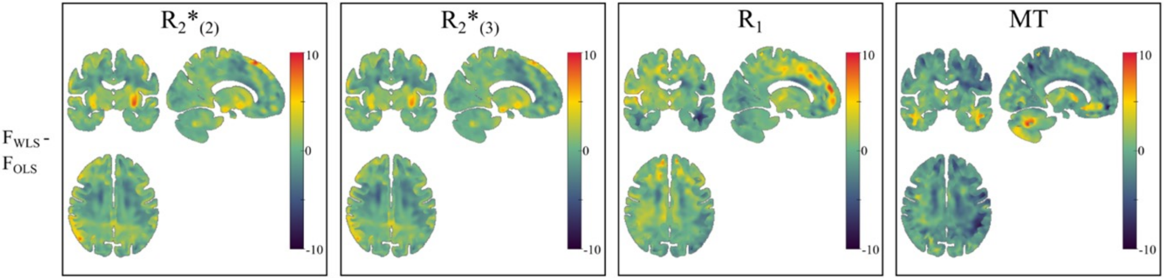


**Figure S1: Noise heteroscedasticity leads to under- or over-estimated statistical results.** Differences in statistical F-maps between OLS and WLS analyses conducted on the full dataset are substantial and may be positive or negative, consistent with the effect of noise heteroscedasticity in OLS analyses.
